# Carboxysome-Inspired Protein Coacervates for Light-Driven CO_2_ Reduction and H_2_ Evolution

**DOI:** 10.64898/2026.09.14.751539

**Authors:** Abesh Banerjee, Roman C Fabry, Giovanna Ghirlanda

## Abstract

Efficient catalysis often requires high local concentrations of reactants and catalysts, which cells achieve through compartmentalization within organelles, such as carboxysomes, that increase the efficiency of bacterial carbon fixation. Here, we engineered a photocatalytic reaction compartment that concentrated an artificial metalloenzyme, carbon dioxide, and a photosensitizer by liquid-liquid phase separation triggered by a cationic polypeptide, deca(L-arginine) (R_10_). At low R_10_ concentrations, CoPPIX binding increases the α-helical structure of the otherwise disordered protein, supercharged cytochrome b_562_(–22). At higher concentrations, electrostatic complexation produces spherical droplets that enrich the protein and cobalt cofactor and recruit the photosensitizer [Ru(bpy)_3_]²⁺. Under illumination, coacervation increased hydrogen evolution 1.9-fold and CO formation from CO_2_ 1.3-fold relative to the corresponding solution-phase protein system. Co-encapsulation of carbonic anhydrase changed the product distribution specifically in the condensed phase: CO production increased 3.2-fold, H_2_ evolution decreased from 1.51 to 0.70 µmol, and CO selectivity among the detected two-electron products rose from 33% to 77%. These results demonstrate that bioinspired coacervates can stabilize reactive intermediates, enrich local substrate concentrations, and integrate multiple catalytic functions, providing a generalizable framework for programmable, light-driven synthetic organelles.

## Introduction

Cells depend on precise spatial and temporal organization to regulate biochemical reactions, sustain metabolic flux, and respond to environmental stimuli.(1–3) Membrane-bound organelles in eukaryotes and protein-based microcompartments in prokaryotes have long been recognized as the principal means of cellular compartmentalization, and a growing body of work has established the importance of membraneless organelles, which are dynamic, liquid-like condensates formed through liquid–liquid phase separation (LLPS).(4–7) These structures arise from multivalent interactions among proteins, nucleic acids, and other biomolecules and are now recognized as key regulators of gene expression, stress response, enzymatic activity, and signaling (8–12). Because they assemble, dissolve, and reorganize rapidly, they offer a means of modulating biochemical functions without lipid boundaries. These properties have motivated bottom-up efforts to construct artificial organelles that localize and control catalytic reactions (13–21).

Complex coacervation provides a particularly useful route to synthetic membraneless compartments, because it is driven by associative phase separation between oppositely charged macromolecules (22–24). Engineered coacervates have been assembled from charged polypeptide pairs, short peptides, globular proteins, protein-polymer complexes, and recombinant protein polymers, and have been used to sequester protein clients, protect enzymes, regulate release, and create protocell-like reaction environments.(25–33) In these systems, phase behavior is controlled not only by net charge, but also by charge density, charge patterning, chain architecture, salt concentration, pH, and stimulus-responsive sequence elements.(21, 34–36) For example, supercharged proteins and ionic peptide tags have shown that presenting charge in a flexible, locally multivalent domain can favor liquid coacervate formation and improve salt tolerance relative to distributing similar charge across a globular surface.(37–41) These studies establish complex coacervation as a programmable strategy for protein recruitment and compartment formation.

However, most engineered protein coacervates treat the folded protein as either a cargo that partitions into a preformed dense phase or as a charged scaffold whose folded state is largely independent of condensation. In contrast, natural biochemical compartments often couple assembly to functional activation, so that structure, localization, and activity emerge together. Here, we sought to build a synthetic condensate in which coacervation is directly linked to protein folding and catalytic output.

Carboxysomes illustrate how compartmentalization improves CO_2_ fixation (42–44). These bacterial microcompartments encapsulate the enzyme RuBisCO and carbonic anhydrase within a protein shell permeable to bicarbonate, elevating local CO_2_ concentrations, suppressing the competing oxygenase reaction, and markedly enhancing carbon fixation efficiency (43, 45). Inspired by this spatial organization, we developed a membraneless coacervate containing a negatively supercharged cytochrome *b*_562_ variant, cobalt protoporphyrin IX (CoPPIX), and the cationic peptide deca(L-arginine) (R_10_). Cobalt-substituted cytochrome *b*_562_ is an established catalyst for light-driven H_2_ evolution and CO_2_ reduction (46, 47), providing a benchmark for measuring the effect of condensation. We show that R_10_ promotes cofactor-dependent folding at low concentration and drives phase separation at higher concentrations. The resulting condensates support light-driven H_2_ and CO production, and recruit carbonic anhydrase to shift product distribution toward CO formation, mirroring its role in natural carboxysomes.

These findings establish a generalizable framework in which supercharged protein engineering, polyelectrolyte-induced coacervation, and artificial metallocofactors operate in concert to create functional, stimuli-responsive condensates. Unlike prior systems that primarily use coacervation to concentrate or deliver folded proteins, this platform couples structural activation of a protein to condensate assembly and catalytic output, enabling the bottom-up construction of synthetic compartments whose formation and function are mechanistically linked.

## Results

### Cofactor-Induced Folding of Supercharged Cytochrome *b*_562_ in the Presence of Polyarginine

Cyt *b*_562_(–22) is an engineered heme-binding protein with a high net negative surface charge. Cyt *b*_562_(–22) is predominantly disordered under physiological buffer conditions because extensive surface supercharging destabilizes its native α-helical fold; addition of monovalent and divalent cations partially compensates for this electrostatic repulsion and permits heme binding, restoring the fold.(48) We reasoned that multivalent cationic peptides such as R_10_ could promote both folding and coacervation in a concentration-dependent manner: at low concentrations, guanidinium side chains could partially neutralize electrostatic repulsion through bidentate hydrogen bonding with surface glutamates, generating a conformational ensemble permissive for cofactor binding.(49) Circular dichroism (CD) spectroscopy showed that apo cyt *b*_562_(–22) remained predominantly unfolded in the presence of 20 µM R_10_, indicating that electrostatic screening by the peptide alone was insufficient to restore the secondary structure under these conditions. Titration of CoPPIX into the cyt *b*_562_(–22)/R_10_ mixture progressively increased the α-helical content, with minima at 208 and 222 nm and saturation near a 1:1 protein-to-cofactor ratio. UV–visible absorption spectroscopy independently supported cofactor binding: the Soret maximum shifted from 417 nm for free CoPPIX to 430 nm after the addition of cyt *b*_562_(– 22), consistent with the spectral signature expected for cyt *b*_562_-bound CoPPIX (refs). Fitting the titration yielded an apparent dissociation constant (K_D,app_) of 0.92 ± 0.03 µM, weaker than the value of 45 ± 19 nM reported for WT cyt *b*_562_.(46) This difference likely reflects the additional energetic cost of folding the supercharged protein, which is thermodynamically coupled to CoPPIX binding (Fig. 2).

**Figure 1.**
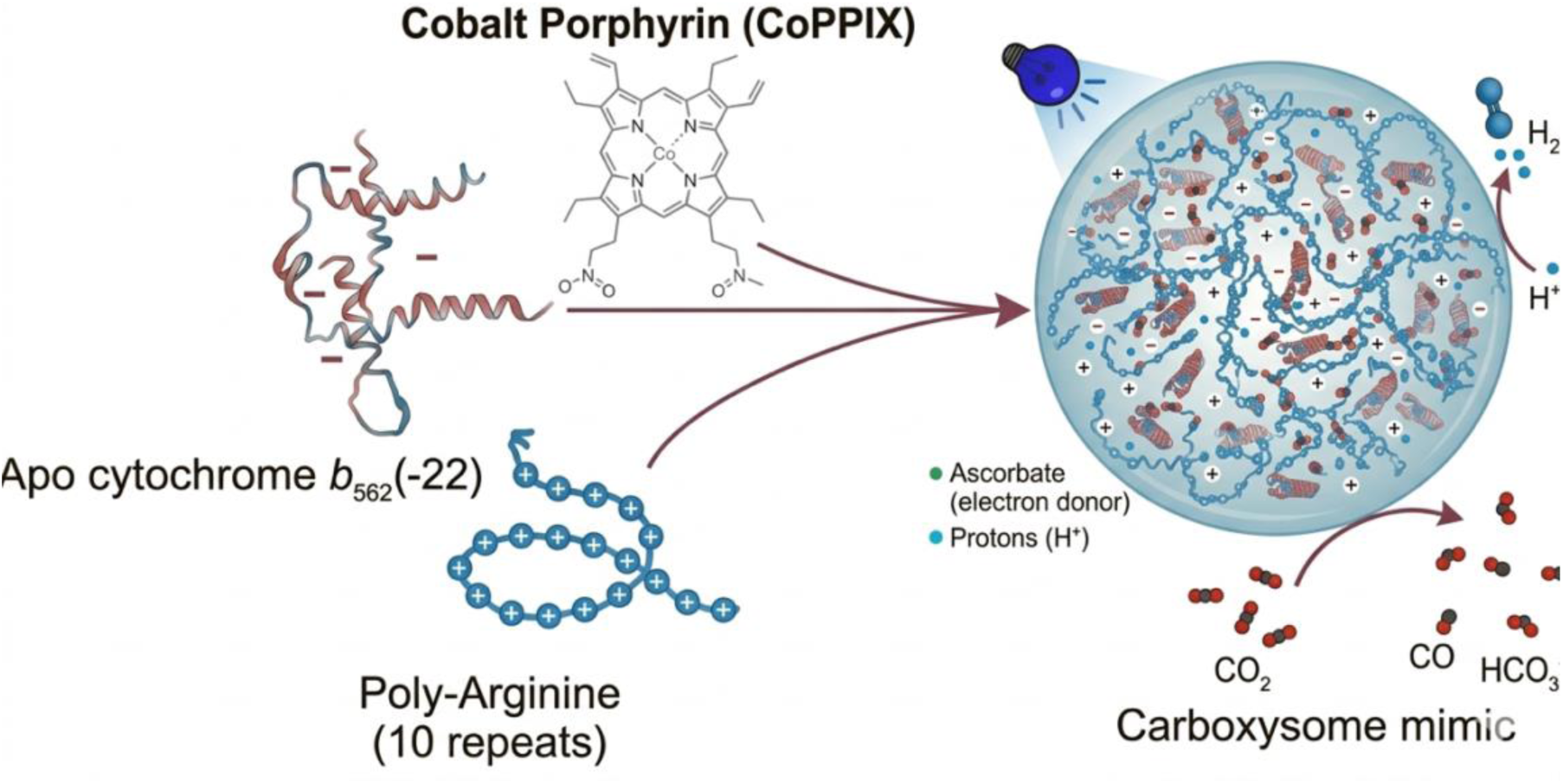
Proposed schematic for the assembly of functional coacervates.

**Figure 2.**
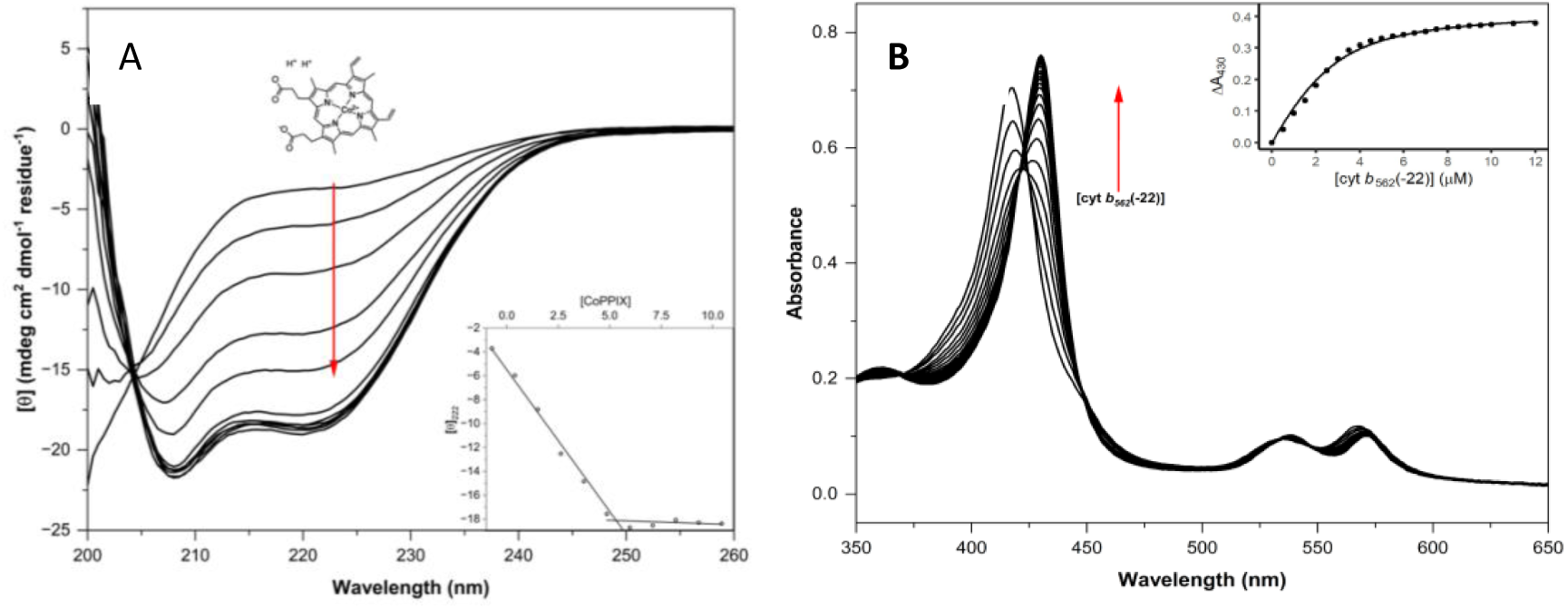
CoPPIX binding induces folding of cyt *b*_562_(–22) at low R_10_ concentrations. (A) Circular dichroism spectra of 20 µM apo-cyt *b_562_*(–22) in 20 µM R_10_ was titrated with increasing concentrations of 4 µM CoPPIX show a progressive gain in α-helical structure, evidenced by the deepening minima near 208 and 222 nm. The red arrow indicates the direction of increasing CoPPIX. Inset: ellipticity at 222 nm plotted as a function of [CoPPIX], demonstrating a 1:1 binding stoichiometry between porphyrin and protein. (B) UV–vis absorption spectra monitoring 0.5 µM cyt *b_562_*(–22) increment titration into 5 µM CoPPIX reveals a bathochromic shift of the Soret band. The red arrow indicates increasing [cyt *b*_562_(–22)]. Inset: tight-binding fit of ΔA₄_3_₀ versus protein concentration. Conditions: (A): cytb562(−22) = 20 µM, 20 µM R_10_, 4 µM increments of CoPPIX, buffer = 20 mM Na-Pi, pH 7.4 (B) 4.865 µM CoPPIX, 2µM R_10_, 0.5 µM increments of cytb562(−22), buffer = 20 mM Na-Pi, pH 7.

### Polyarginine drives complex coacervation

We tested whether higher R_10_ concentrations drive complex coacervation with folded cyt *b*_562_(–22). Turbidity at 650 nm was measured for CoPPIX– cyt *b*_562_(–22)/R_10_ mixtures at a total macromolecule concentration of 1 mg mL⁻¹ in 20 mM sodium phosphate, pH 7.4. The resulting turbidity profiles, plotted against the positive charge fraction (f⁺), showed a bell-shaped dependence characteristic of complex coacervation (Fig. S2). Turbidity was maximal at f⁺ ≈ 0.8 and decreased at both low f⁺ (protein-rich) and high f⁺ (R_10_-rich) compositions, where excess like charge disfavors condensate formation.(50) Apo cyt *b*_562_(–22) /R_10_ mixtures behaved similarly; subsequent coacervate experiments were carried out at f+ = 0.76, at which confocal fluorescence microscopy showed spherical droplets. In CoPPIX–cyt *b*_562_(–22)/R_10_ coacervates, C343-labeled cyt *b*_562_(–22) and CoPPIX colocalized within the droplets (Fig. 3A-C). Fluorescence intensities yielded partition coefficients of 46 ± 1 for C343-labeled cyt *b*_562_(–22) and 63 ± 1 for CoPPIX (Table S1), demonstrating enrichment of both components in the dense phase.

**Figure 3.**
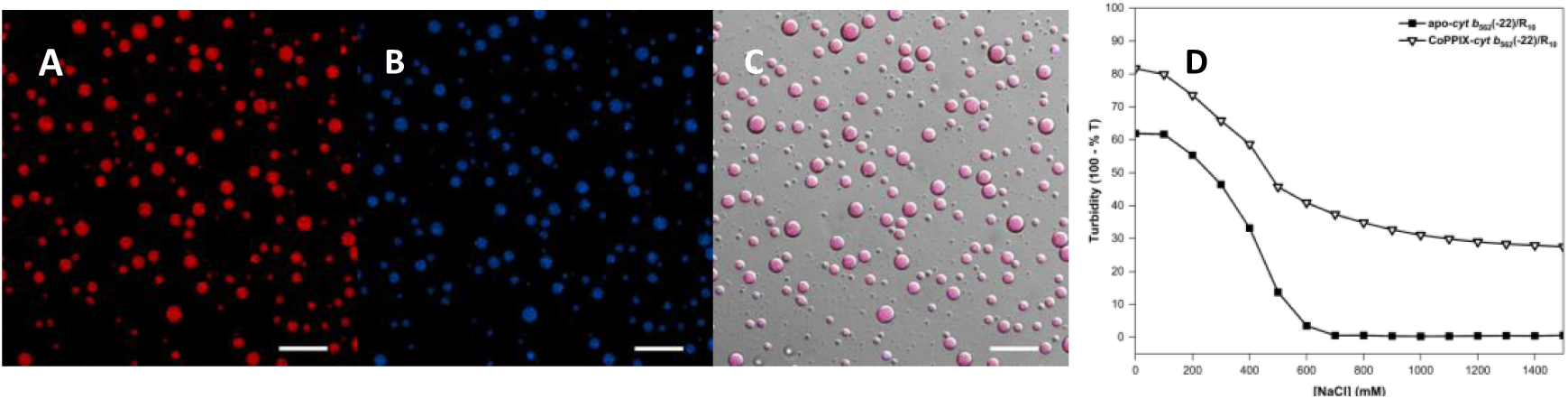
Complex coacervation behavior of CoPPIX–cyt b_562_(–22)/R_10_ at optimal coacervation conditions ( 40 uM CoPPIX-cytb562(−22), 273.6 uM R10, buffer = 20 mM Na-Pi, pH 7.4, f+ = 0.76). Confocal fluorescence and brightfield images. (A) CoPPIX fluorescence (red). (B) C343-labeled cyt *b_562_*(–22) (blue). (C) merged brightfield/fluorescence image, illustrating spherical, liquid-like droplets enriched in both protein and cofactor. Scale bars: 20 µm. (D) Salt titration profiles for apo-cyt *b_562_*(–22)/R_10_ and CoPPIX–cyt *b_562_*(–22)/R_10_, showing changes in turbidity with increasing salt concentration.

### Salt Sensitivity of the Coacervates

Increasing NaCl concentration progressively reduced turbidity in both apo and CoPPIX-bound mixtures (Fig. 3D), supporting a major contribution from electrostatic interactions as a driving force for coacervation. Fitting the transitions yielded apparent critical salt concentrations of 413 mM and 435 mM for the apo and CoPPIX-bound mixtures, respectively. Apo samples returned to baseline turbidity above approximately 700 mM NaCl, whereas CoPPIX-bound samples retained approximately one-third of their initial turbidity at 1.5 M NaCl.(51)

### Folding occurs in the condensed phase

To determine whether apo and CoPPIX–cyt *b*_562_(–22) were folded within the coacervates, we examined their secondary structure and cofactor environment using circular dichroism (CD) and UV–visible spectroscopy. Spectra were collected on droplet suspensions. in a 0.1 mm cuvette to reduce attenuation by scattering, and a matched R_10_-only sample was subtracted. Both apo and CoPPIX–cyt *b*_562_(–22) displayed minima at 208 and 222 nm, consistent with α-helical structure. Apo cyt b_562_(–22), which is largely disordered in dilute solution at the R_10_ concentration used in Fig. 2A, produced a substantial helical signal in the coacervate suspension ([θ]_222_ = 19,400 mdeg cm² dmol⁻¹ residue⁻¹, approximately 42.3% helix), indicating that partitioning into the coacervate is sufficient to drive partial folding. The CoPPIX-bound cyt *b*_562_(–22) produced a larger negative ellipticity at 222 nm, consistent with higher helical content ([θ]_222_ = 25,000 deg cm^2^ dmol^-1^; approximately 56.4% helix). This behavior mirrors the difference in helical content observed for apo and Co-PPIX bound WT cyt *b*_562_.(46, 47)

The UV–visible spectrum of CoPPIX–cyt *b*_562_(–22) in the droplet suspension showed a Soret maximum at 427 nm (Fig. 4B), consistent with protein-bound CoPPIX and similar to the reported spectrum of Co-substituted WT cyt *b*_562_.(46, 47) The spectrum showed no obvious broadening or blue-shifted shoulder associated with extensive porphyrin self-association.

**Figure 4.**
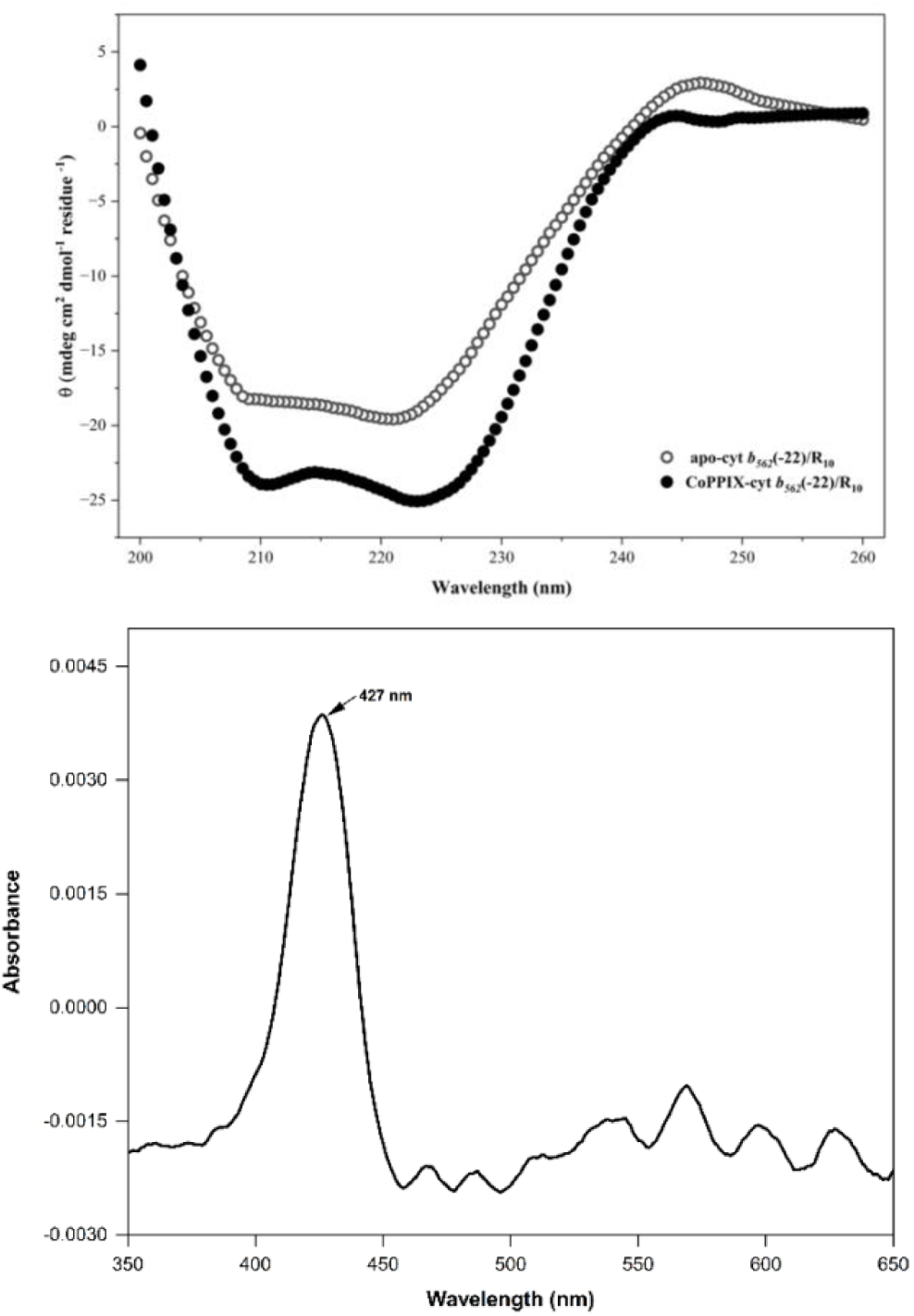
Secondary structure and cofactor environment of cyt *b*_562_(–22) within the coacervates. (A) CD spectra of apo cyt *b*_562_(–22)/R_10_ and CoPPIX–cyt *b*_562_(–22)/R_10_ coacervates. Both display the 208/222 nm minima of the α-helical structure; CoPPIX incorporation increases the magnitude of the negative ellipticity, indicating a higher helical content. The spectra were corrected for the R_10_ contribution by subtracting a matched peptide-only sample. (B) UV–visible absorption spectrum of CoPPIX–cyt *b*_562_(–22)/R_10_ droplets, showing a Soret maximum at 427 nm, consistent with the maximum of Co-substituted WT cyt *b*_562_ and with the intact His/Met ligation. Conditions: 20 mM NaPi, pH 7.4, *f*⁺ = 0.76; 40 µM protein, 274 µM R_10_, 0.1 mm path length, 25 °C; *n* = 3 independent preparations.

Together, the data support a model in which R_10_ mediated condensation promote chain compaction of the apo cyt b_562_(–22) to a molten globule-like state, and cofactor binding further shifts the conformation to the folded state.

### Photocatalytic Activity of CoPPIX–cyt *b*_562_(–22) Coacervates

Cobalt-substituted WT cyt *b*_562_ supports photocatalytic hydrogen evolution and carbon dioxide reduction to CO in the presence of a photosensitizer, [Ru(bpy)_3_]²⁺, and a sacrificial electron donor, ascorbate.(46, 47) We therefore tested whether condensation alters the catalytic activity of CoPPIX–cyt b_562_(–22). In the reductive quenching pathway proposed for these systems, photoexcited [Ru(bpy)_3_]²⁺ is reduced by ascorbate to generate a reactive Co(I) species that can support both hydrogen evolution and CO_2_ reduction (refs).

We compared H_2_ production by free CoPPIX, solution-phase CoPPIX–cyt b_562_(–22), and CoPPIX– cyt b_562_(–22)/R_10_ coacervates under argon. After 2 h of illumination, the samples produced 0.60 ± 0.04, 1.72 ± 0.13, and 3.18 ± 0.18 µmol H_2_, respectively (Fig. 5A). Incorporation into the protein increased H_2_ production 2.9-fold relative to free CoPPIX, and condensation produced a further 1.9-fold increase, corresponding to an overall 5.3-fold enhancement relative to free CoPPIX.

**Figure 5.**
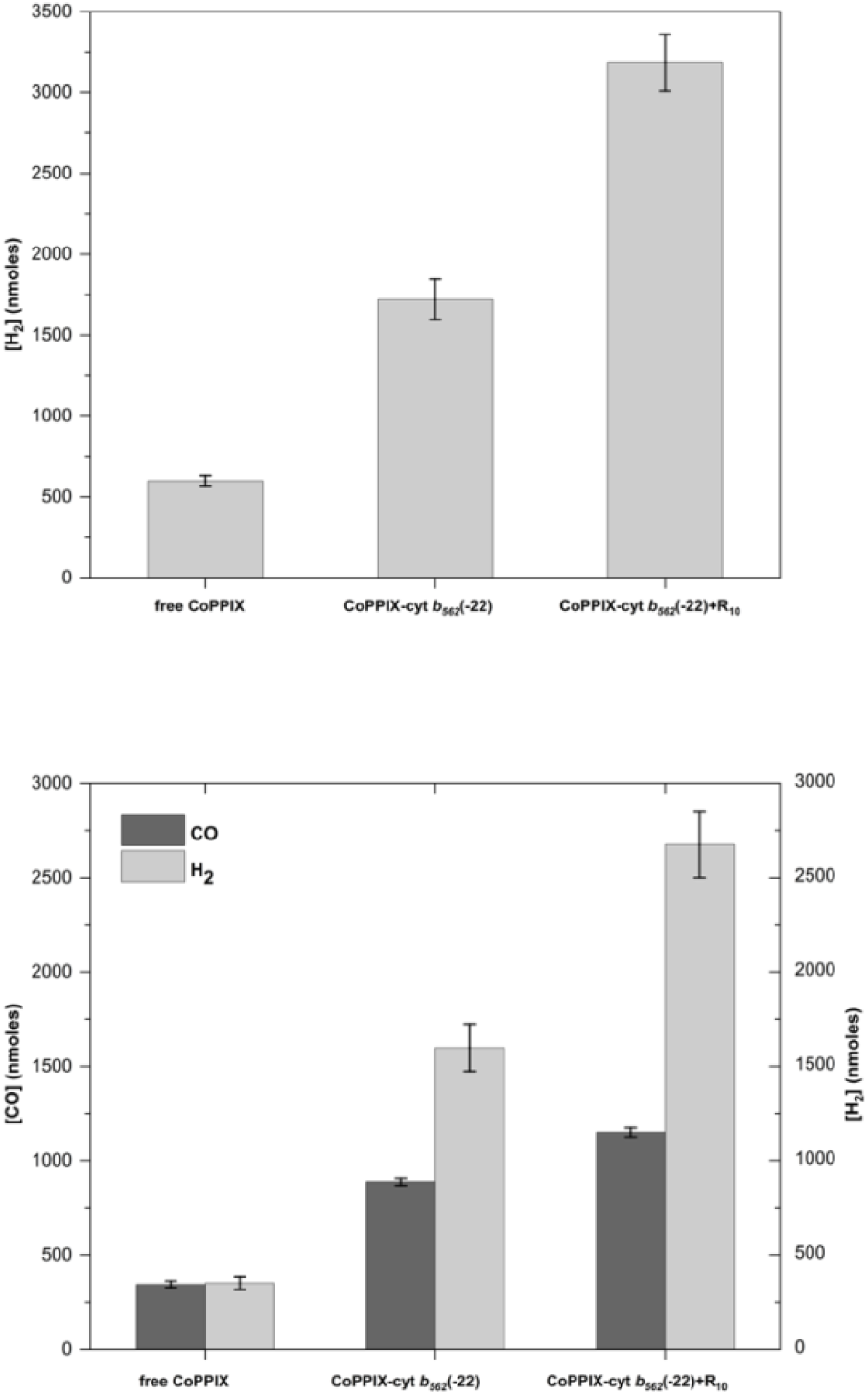
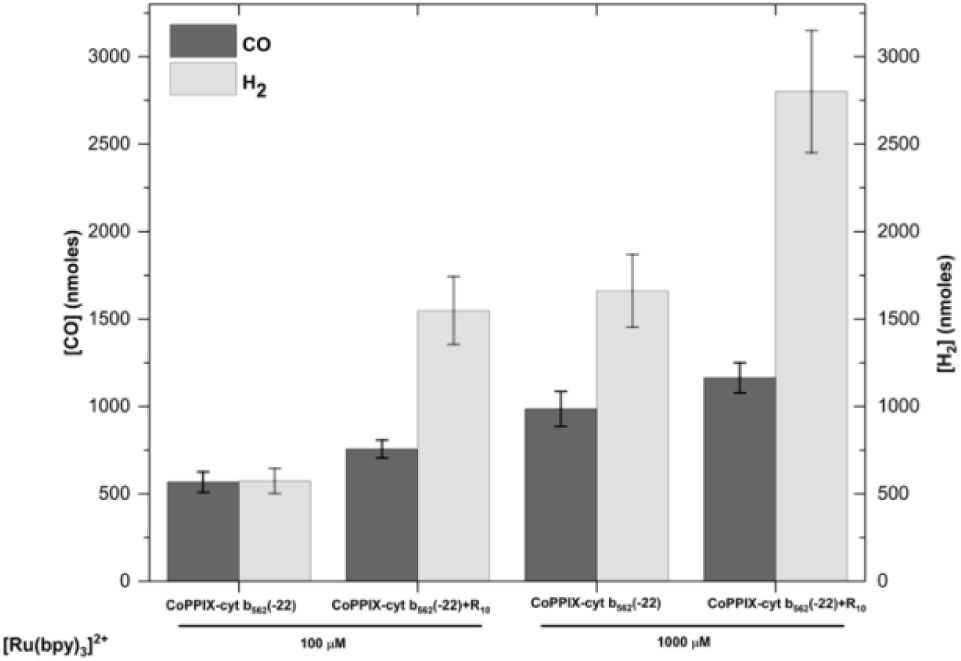
Coacervate encapsulation enhances light-driven H_2_ and CO production by CoPPIX–cyt *b*_562_(– 22). (A) H_2_ production by free CoPPIX, solution-phase CoPPIX–cyt *b*_562_(–22), and CoPPIX–cyt *b*_562_(–22)/R_10_ coacervates under an argon atmosphere. (B) CO (dark bars) and H_2_ (light bars) formed under a CO_2_ atmosphere; CoPPIX–cyt *b*_562_(–22)/R_10_ coacervates are more active than solution-phase CoPPIX–cyt *b*_562_(– 22) for both products. (C) CO (dark bars) and H_2_ (light bars) produced by solution-phase and coacervate samples at two concentrations of [Ru(bpy)_3_]²⁺ (100 and 1000 µM). Conditions: 40 µM protein, 8 µM CoPPIX, 150 mM ascorbate, 1000 µM [Ru(bpy)_3_]²⁺, 100 mM Na-Pi, pH 7.4, f⁺ =0.76, 450 nm illumination for 2 h at 25°C. Error bars represent the standard deviation from n = 4 independent replicates.

Under a carbon dioxide atmosphere, CO formation occurred alongside hydrogen evolution (Fig. 5B). Free CoPPIX produced 0.35 ± 0.02 µmol CO and 0.35 ± 0.04 µmol H_2_; the solution-phase holo protein produced 0.89 ± 0.02 µmol CO and 1.60 ± 0.13 µmol H_2_; and the coacervate produced 1.15 ± 0.03 µmol CO and 2.68 ± 0.18 µmol H_2_. Relative to the solution phase protein, coacervation increased CO formation 1.3-fold and H_2_ evolution 1.7-fold. The CO:H_2_ ratio decreased from approximately 1.0 for free CoPPIX to 0.55 for the solution-phase protein and 0.43 for the coacervate, indicating that both the protein scaffold and the condensed phase increased hydrogen evolution more strongly than CO formation under these conditions.

To test whether photosensitizer recruitment contributes to the catalytic enhancement, we examined the distribution of [Ru(bpy)_3_]²⁺ and varied its bulk concentration. Confocal fluorescence microscopy showed that [Ru(bpy)_3_]²⁺ colocalized with apo cyt b_562_(–22)/R_10_ droplets (Fig. S3); holo-protein condensates could not be resolved in this channel because [Ru(bpy)_3_]²⁺ emission overlapped with CoPPIX emission. The catalytic advantage of the coacervate was greatest at the lower photosensitizer concentration (Fig. 5C). At 100 µM [Ru(bpy)_3_]²⁺, the coacervate produced 1.3-fold more CO and 2.7-fold more H_2_ than the solution-phase protein, whereas at 1000 µM the corresponding enhancements were 1.3- and 1.7-fold. These observations are consistent with a contribution from photosensitizer recruitment.

### Effect of Carbonic Anhydrase on Photocatalysis

In natural carboxysomes, carbonic anhydrase (CA) is co-encapsulated with RuBisCO, where it accelerates the interconversion of bicarbonate and CO_2_, thereby sustaining the supply of CO_2_ to the carboxylating enzyme (16). We investigated whether co-encapsulating CA within our synthetic condensates would similarly influence CO_2_ reduction and the partitioning of electrons between proton reduction (H_2_) and CO_2_ reduction (CO). Confocal imaging confirmed that Cy3-labeled CA partitioned efficiently into the condensates and colocalized with C343-labeled CoPPIX–cyt *b*_562_(–22) (Figure S4).

Under a CO_2_ atmosphere, co-encapsulation of CA markedly changed the product distribution (Fig. 6). CO production increased 3.2-fold, from 0.74 ± 0.14 to 2.40 ± 0.09 µmol, whereas H_2_ production decreased from 1.51 ± 0.06 to 0.70 ± 0.03 µmol. CO selectivity among the detected two-electron products therefore increased from 33% to 77%, and their combined amount increased 1.4-fold, from 2.25 to 3.10 µmol. This redistribution was specific to the condensed phase under the tested conditions. In a homogeneous solution, CA increased CO formation only 1.2-fold, from 0.55 ± 0.05 to 0.66 ± 0.07 µmol, and did not measurably change H_2_ production (0.61 ± 0.03 versus 0.64 ± 0.03 µmol). CO selectivity changed only from 47% to 51%. Note that the carbonic-anhydrase experiments were carried out at 100 µM [Ru(bpy)_3_]²⁺ (the photosensitizer-limiting regime of Fig. 5C); absolute product amounts are therefore lower than in Fig. 5B, which used 1 mM [Ru(bpy)_3_]²⁺.

**Figure 6.**
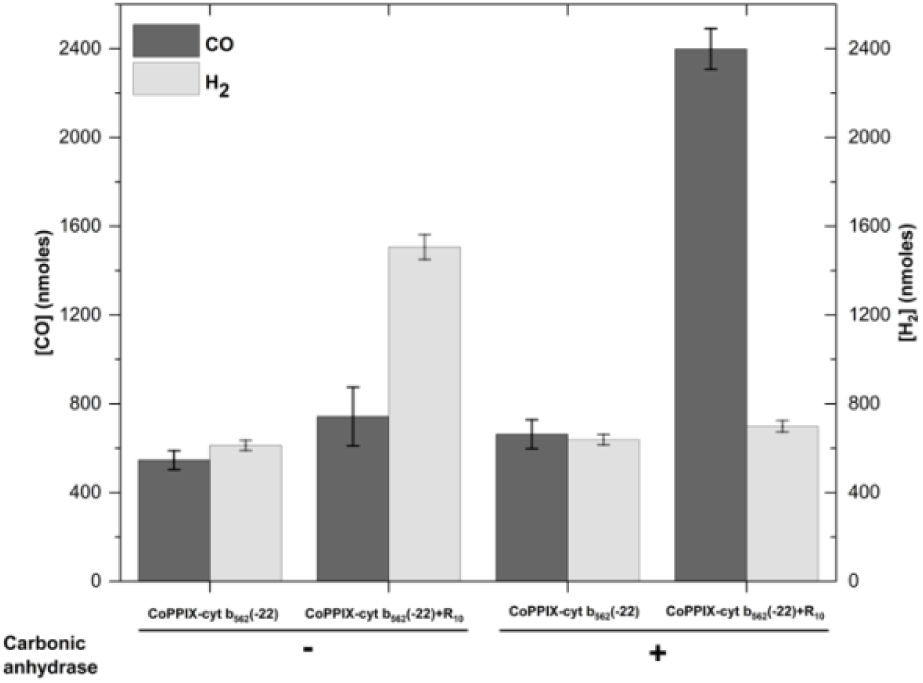
Carbonic anhydrase redirects photocatalysis toward CO_2_ reduction in coacervates. CO (dark bars) and H_2_ (light bars) produced by CoPPIX–cyt *b*_562_(–22) in solution and in R_10_ coacervates, in the absence (left) or presence (right) of carbonic anhydrase under a CO_2_ atmosphere. Within the coacervate, CA increases CO formation by 3.2-fold while decreasing H_2_ evolution by more than half, and the solution-phase control is largely unaffected. Conditions: 40 µM *cytb_562_*(−22), 8 µM CoPPIX, 2 µM carbonic anhydrase, 100 µM [Ru(bpy)_3_]²⁺, 150 mM ascorbic acid, 100 mM Na-Pi, pH 7.4, f⁺ = 0.76, and 450 nm illumination for 2 h. Error bars represent the standard deviation from n = 4 independent replicates.

CA catalyzes CO_2_–bicarbonate interconversion without changing the equilibrium position. The observed change in product distribution may therefore reflect altered reaction kinetics or physicochemical conditions within the condensed phase rather than a higher equilibrium CO_2_ concentration. The reciprocal change in CO and H_2_ production is consistent with competition between CO_2_ and proton reduction at a reduced cobalt intermediate.

## Conclusions

We have shown that negatively supercharged cyt b_562_(–22) links cofactor-dependent folding to complex coacervation with R_10_. At low R_10_ concentration, CoPPIX binding increases α-helical structure. At higher R_10_ concentration, the protein and cofactor partition strongly into spherical coacervate droplets, where spectroscopy remains consistent with protein-bound CoPPIX and increased protein helicity.

The coacervates also enhance photocatalysis. Relative to free CoPPIX, encapsulation increased hydrogen evolution 5.3-fold and CO_2_ reduction to CO 3.3-fold; relative to the solution-phase holo-protein, the gains were 1.9- and 1.3-fold. The advantage was greatest when the photosensitizer was limiting and narrowed at high [Ru(bpy)_3_]²⁺, implying that the co-partitioning of the photosensitizer with the catalyst was the principal origin of the effect rather than a general medium effect. Condensation alone favors proton reduction over CO_2_ reduction.

However, co-encapsulating carbonic anhydrase switched the product distribution: CO formation increased 3.2-fold, whereas H_2_ evolution decreased by more than half, raising the CO share of the two-electron product from 33% to 77%.

These results define a design strategy for functional coacervates in which surface-charge engineering programs both the activation of a metalloprotein and its assembly into a functional compartment, and a second enzyme, recruited into that compartment, redirects catalytic selectivity.

## Acknowledgments

We thank Prof. Ronald Koder (CUNY) for gifting the plasmid coding for cyt*b_562_*(−22). This work was supported by NSF grants 1935105 to GG.

## Supplementary information

## Materials and Methods

### Protein Expression and Purification

E. coli BL21(DE3) cells were transformed with a pET32a(+) (Novagen) plasmid encoding for cyt *b_562_*(−22), gifted by R. Koder (CUNY), and plated onto agar plates containing ampicillin. Single colonies were picked and used for 50 ml starter cultures, which were inoculated into 2×YT–ampicillin medium and grown at 37 °C until OD₆₀₀ ≈ 0.6. Expression was induced with 1 mM IPTG and continued for 6 h at 37 °C. Cells were harvested and resuspended in lysis buffer (50 mM NaPi, pH 8.0, 300 mM NaCl, 10 mM imidazole), then lysed by ultrasonication (30 s on/30 s off duty cycle, 20 min total processing time, on ice). The lysate was clarified by centrifugation and purified by Ni–NTA affinity chromatography using a 5 mL HisTrap HP column (Cytiva, Marlborough, MA, USA) connected to an ÄKTA Pure chromatography system (Cytiva, Marlborough, MA, USA). Bound protein was eluted through a 20–500 mM imidazole gradient, with the target eluting near 400 mM imidazole.

Thioredoxin A fusion tags were removed by TEV protease cleavage (80:1 protein to TEV ratio) overnight at 4 °C in TEV buffer (50 mM NaPi, pH 8.0, 1 mM DTT, 0.5 mM EDTA). The target protein was isolated from the reaction mixture by Ni–NTA affinity chromatography as flow-through fraction. The final purification was carried out via reverse-phase HPLC using a C18 column (Phenomenex, Torrance, CA). The protein mass was confirmed by MALDI-TOF MS on a Bruker RapifleX MALDI-TOF/TOF instrument (Bruker Corporation, Billerica, MA, USA) operating in linear positive-ion mode; samples were desalted with C18 ZipTips (MilliporeSigma) and co-crystallized with an α-cyano-4-hydroxycinnamic acid matrix.

### Cofactor-Binding Assays

20 µM of cyt *b_562_*(−22) was prepared in 20 mM NaPi, pH 7.4, containing 20 µM R_10_. CoPPIX stocks (500 µM in 1 M KOH) were diluted into the buffer. CD spectra (200–260 nm) were collected in a 1 mm quartz cuvette at 25 °C on a Jasco 815 instrument (Jasco Inc. Easton, MD), equipped with a Peltier thermoelectric temperature controller. CoPPIX was titrated into the apo-protein in 4 µM increments, equilibrated for 15 minutes, and three consecutive scans were averaged. Folding transitions were monitored by observing the development of α-helical minima at 208 and 222 nm. UV-visible spectroscopic monitoring of CoPPIX binding was carried out on a Cary 50 spectrophotometer(Agilent, Santa Clara, CA): CoPPIX (approximately 5 µM) was prepared in 20 mM NaPi, pH 7.4, containing 20 µM R_10_ in a 1 cm cuvette. Apo-protein was titrated in 0.5 µM increments with 15-minute equilibration between additions. The Soret band at 427 nm was monitored to assess binding. Apparent dissociation constants (K*_D_,_app_*) were determined by analyzing ΔA versus total protein concentration using a tight-binding model equation as described previously.(Alcala-Torano et al., 2021; Sommer et al., 2016)

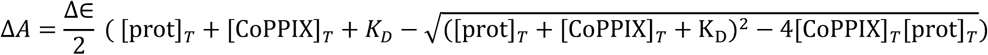

Where ΔA and Δɛ are the differences in absorbance and extinction coefficients, respectively, between the free porphyrin and the holo protein at the latter’s Soret peak. [CoPPIX]_T_ is the total CoPPIX concentration, and [prot]_T_ is the total protein concentration after each addition.

### Turbidity Assays for Coacervation

Apo-*cytochrome b_562_*(–22) was dissolved in 20mM Na-Pi, 20 µM R10, pH 7.4; CoPPIX was prepared as a 5.8 mM stock in 1 M KOH. Equal concentrations of apo-protein and CoPPIX (82 μM; 1 mg/mL) were combined in 20 mM Na-Pi, 20 µM R10, pH 7.4, and incubated for 30 min to allow holoprotein formation. Polyarginine (R10) was prepared as a 526 µM (1 mg/mL) stock in 20 mM Na-Pi, pH 7.4. Holoprotein and R10 were combined at protein mass fractions of 0–1 (0.04 increments, corresponding to varying charge fractions) in a 96-well glass-bottom plate (Cellvis P96-1.5H-N); absorbance at 650 nm was used to monitor phase separation. Charge fractions were calculated from mass fractions, and %T was calculated from absorbance (A) as A = −log(%T); turbidity (100 − %T) was plotted against positive charge fraction to generate turbidity profiles. To directly relate macromolecular composition to electrostatic stoichiometry, the mass fraction of protein was converted to positive charge fraction using:

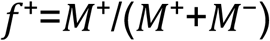

where M-is the total negative charge (calculated from protein mass divided by the molar mass of the protein, multiplied by its net negative charge), and M+ is the total positive charge (calculated from protein mass divided by the molar mass of the protein, multiplied by the net positive charge per monomer).

### Fluorescent Labeling of Proteins

C343 NHS ester dye (ThermoFisher Scientific) was added to apo cyt *b_562_*(−22) at a 10:1 dye to protein molar ratio in 20 mM NaPi at pH 8.0. Reactions continued overnight at 4 °C. Excess dye was removed using PD-10 desalting columns (Cytiva, Marlborough, MA, USA; Sephadex G-25 resin). Cy3 NHS ester (Lumiprobe) labeling of carbonic anhydrase (C3934, Sigma-Aldrich, Inc) followed the same procedure.

### Confocal Microscopy of Coacervates

Cyt *b_562_*(−22)–R_10_ mixtures were prepared at compositions matching maximum turbidity and placed onto PEG-silane–treated slides with silicone isolators (coverslips were cleaned in KOH/isopropanol and detergent, then PEG-silane–functionalized with 5 mg mL⁻¹ PEG-silane in 96% ethanol containing 1% (v/v) glacial acetic acid at 70 °C for 30 min; silicone isolators from Thermo Fisher Scientific). Samples were imaged within 30 minutes using excitation/emission of 405/590– 620 nm (CoPPIX), 445/460–490 nm (C343), and 561/565–575 nm (Cy3). Partition coefficients (PCs) were determined by dividing the mean fluorescence intensity inside droplets by that of the surrounding dilute phase. Imaging was performed on a Nikon A1R HD25 laser-scanning confocal microscope (Nikon Instruments Inc., Melville, NY, USA) equipped with a Plan Apo VC 60× water-immersion objective (NA 1.2) in Galvano scanning mode; images were processed in FIJI/ImageJ.

### Spectroscopy of Proteins in Coacervates

The cyt *b_5C2_*(−22)-R_10_ coacervates (40 µL) were formed directly in a 0.1 mm quartz cuvette (demountable 0.1 mm path length quartz cell, 20/C-Q-0.1, Starna Cells Inc., Atascadero, CA, USA). A 40 µL phase-separated sample was formed directly in the cuvette and incubated for 30 min at room temperature before sealing. CD spectra were collected from 260–200 nm at 50 nm/s with a 4 s integration time at 25 °C, averaging six scans; an equivalent spectrum of 273.6 µM R10 (the concentration used in the coacervates) was collected identically and subtracted from the coacervate spectrum to isolate the holoprotein CD signal. UV-vis spectra of the coacervate were likewise collected using the same 0.1 mm cuvette.

### Photocatalytic studies

Photocatalytic H_2_ and CO_2_-reduction assays were performed using CoPPIX–cyt b_562_(–22) as catalyst, [Ru(bpy)_3_]^2+^ as photosensitizer (Sigma-Aldrich, Inc), and ascorbic acid (Sigma-Aldrich, Inc) as sacrificial electron donor, with and without R10 coacervation. CoPPIX (dissolved in 1 M KOH, diluted 1:1000 into 500 mM Na-Pi, pH 7.4) and cyt b_562_(–22) concentrations were determined by UV-vis, and CoPPIX was incubated with a 5-fold excess of cyt b_562_(–22) for 30 min at room temperature to form the holoenzyme. Non-coacervate reactions (100 mM Na-Pi, pH 7.4) and coacervate reactions (100 mM Na-Pi, pH 7.4, plus 273.6 µM R10) contained final concentrations of 40 µM cyt b_562_(–22), 8 µM CoPPIX, 150 mM ascorbic acid, and 1 mM [Ru(bpy)_3_]^2+^ in a total volume of 500 µL in 2 mL glass vials fitted with spin vanes and sealed with crimp tops. For CO_2_-reduction assays, buffers were pre-saturated with CO_2_ (2 h bubbling, pH readjusted) in place of the standard buffers; reaction assembly was otherwise identical. Vials were purged with argon (H_2_ assay) or CO_2_ (CO_2_-reduction assay) for 30 min via needle through the septum, with a control vial confirming minimal residual O_2_ by GC, then illuminated for 2 h at 25 °C with stirring in a water-circulated photocatalytic chamber (HepatoChem Temperature Controlled PhotoRedOx Box) with 450 nm illumination (EvoluChem 55 W/cm^2^ blue LED). Vials were inverted and rested in the dark for 15 min; 100 µL of argon was injected via a Hamilton gas-tight syringe, and 100 µL of headspace gas was withdrawn and analyzed by gas chromatography (GC) (SRI 310 C, SRI Instruments, Torrance, CA). H_2_ and CO were quantified simultaneously from a single injection using a thermal conductivity detector (TCD) and a flame ionization detector (FID) with methanizer connected in series, with argon carrier gas and a temperature program of 40 °C (4 min hold) ramped at 50 °C/min to 180 °C (held to CO_2_ elution, ∼10 min; total CO_2_ retention ∼16 min); H_2_ eluted on the TCD at 0.400 min and CO on the FID at ∼3.42 min. Peak areas were converted to moles using H_2_ and CO calibration curves (SI Appendix); all GC measurements were performed in triplicate unless stated otherwise.

To assess carbonic anhydrase (CA) localization, coacervates formed from CoPPIX–cyt b_562_(–22) (5% C343-labeled) and R10, at f^+^ = 0.76, were mixed with 2 µM Cy3-labeled CA, applied to a PEGylated glass slide, and imaged on a Nikon AX R laser scanning confocal microscope (Ti2-E, 60×/1.42 NA oil objective) using 405, 445, and 561 nm excitation for CoPPIX, C343, and Cy3, respectively. Photocatalysis with and without CA was compared using coacervate reactions (100 mM Na-Pi, CO_2_-saturated) containing 40 µM cyt b_562_(–22), 8 µM CoPPIX, 150 mM ascorbic acid, 100 µM [Ru(bpy)_3_]^2+^, and, where indicated, 2 µM CA, assembled and analyzed as described above.

---

**Figure S1.**
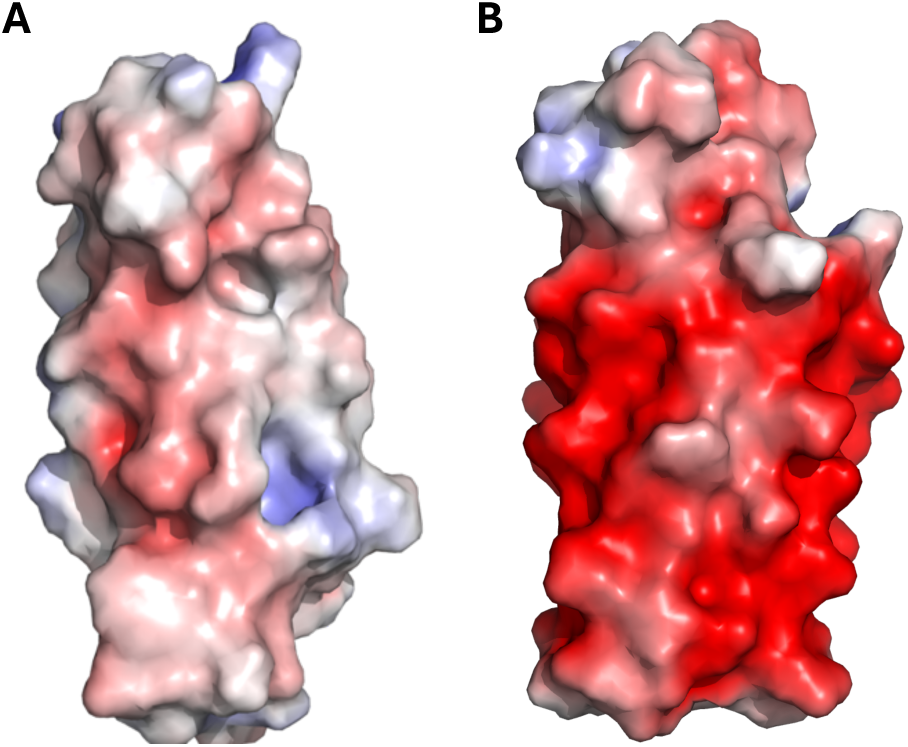
Electrostatic surface visualization of cyt *b*_562_ variants highlights charge redistribution upon supercharging. Surface representations show the native protein (left) and the supercharged cyt *b*_562_(–22) variant (right), with electrostatic potentials mapped from negative (red) to positive (blue). Introduction of acidic residues dramatically increases the overall negative surface charge.

**Figure S2.**
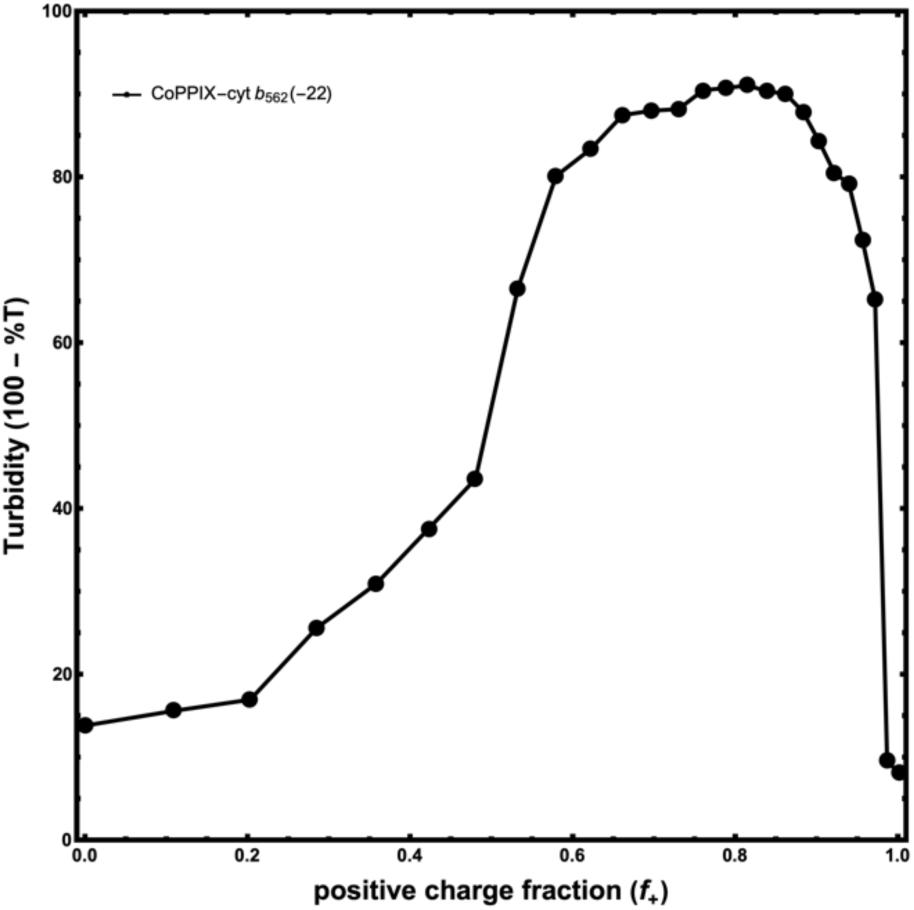
Turbidity screening of Co-cyt *b562*(−22) and R10; total macromolecular concentration 1 mg/mL in 20 mM Na-Pi, pH 7.4. Turbidity (A₆₅₀) is plotted as a function of charge fraction f⁺.

**Figure S3:**
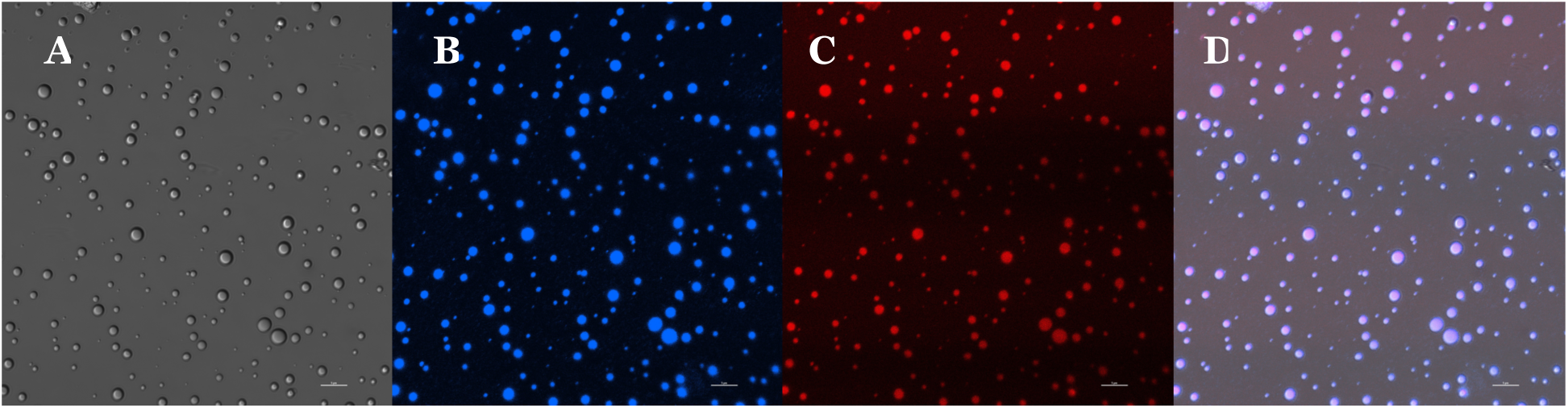
Colocalization of [Ru(bpy)_3_]²⁺ within cyt b_562_(–22)/R_10_ coacervates. (A) Brightfield image of droplets formed under optimal LLPS conditions. (B) C343-labeled cyt b_562_(−22) (Ex: 445 nm; Em: 460-490 nm) enriches strongly within the condensed phase. (C) [Ru(bpy)_3_]^2+^ fluorescence (Ex: 405 nm; Em:590-620 nm) shows photosensitizer co-localization. (D) Merged brightfield/fluorescence image confirms co-recruitment of protein and photosensitizer into the same droplets. Scale = 20 μm. Conditions: 20 mM NaPi, pH 7.4, f* = 0.76, 40 μM apo-cyt b_562_(−22) (with 5% C343 labeled apo-cyt b_562_(−22)), 273.60 μM R, 2 μM [Ru(bpy)_3_]^2+^

**Figure S4.**
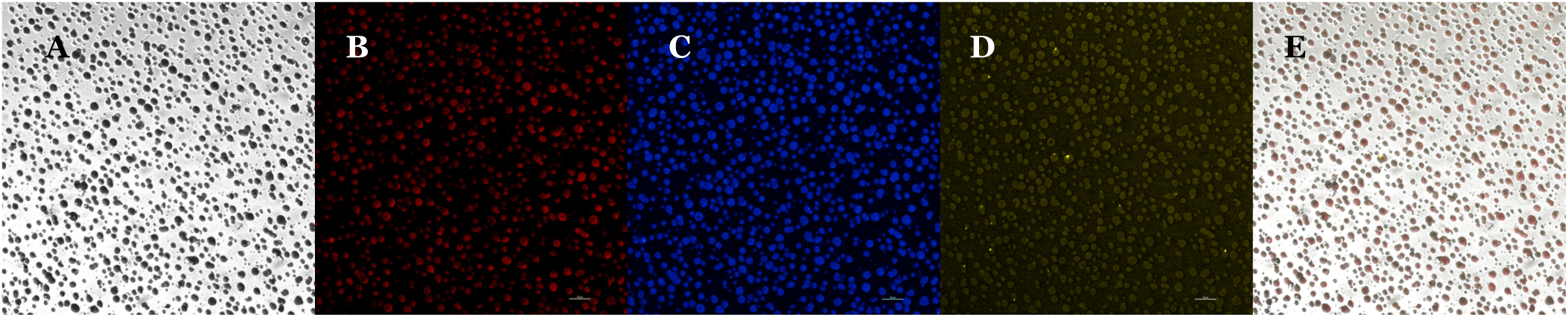
Recruitment of carbonic anhydrase into CoPPIX–cyt b_562_(–22)/R_10_ coacervates. (A) Brightfield image. (B) CoPPIX fluorescence (red). (C) C343-labeled cyt b_562_(–22) fluorescence (blue). (D) Cy3-labeled carbonic anhydrase fluorescence (yellow). (E) Merged brightfield and fluorescence image. Scale bars, 20 µm.

**Figure S5.**
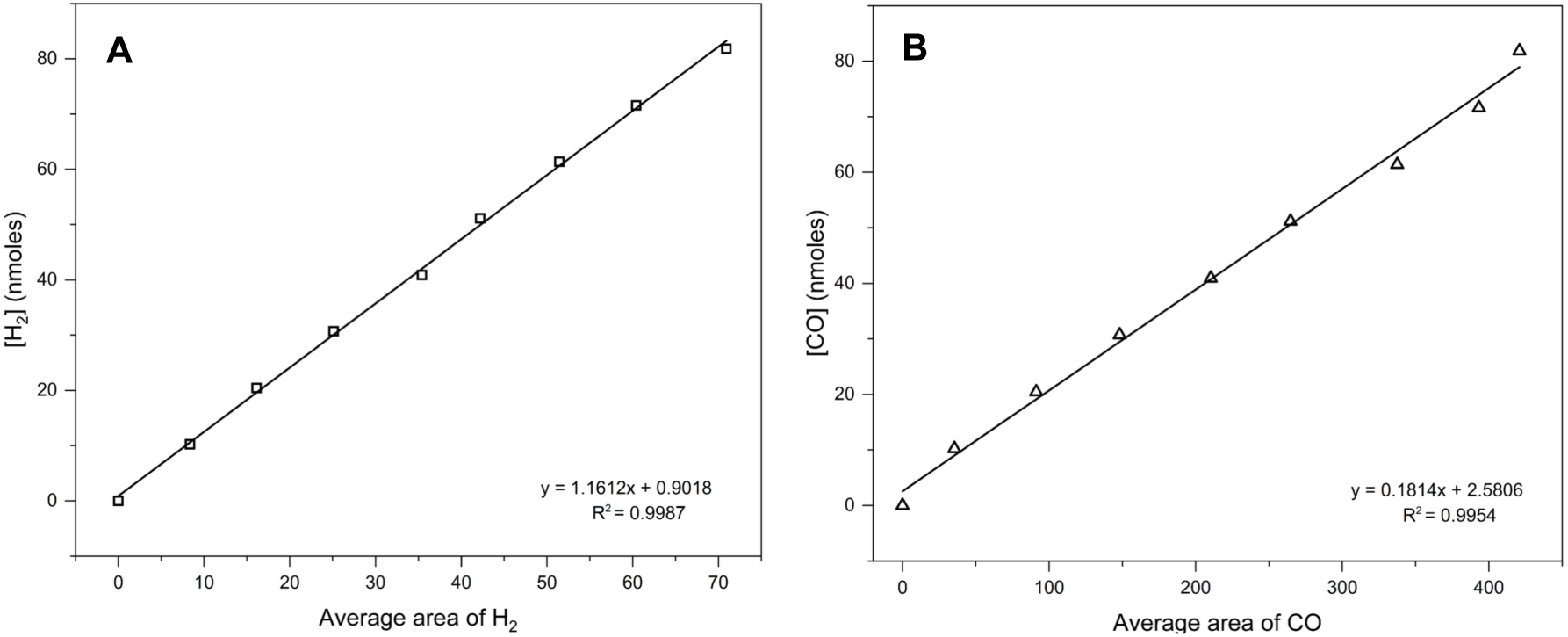
GC calibration curves. Corresponding calibration curve for H_2_ **(A)** and CO **(B)**. These standards were used to convert GC peak areas into nanomoles of product for all catalytic assays.

**Table S1.** Partition coefficients of the supercharged protein and the CoPPIX in the CoPPIX-supercharged protein/R_10_ coacervates, obtained by analyzing confocal microscopy images using ImageJ; n=60 (Figure S4)

|  |  |  |
| --- | --- | --- |
| CoPPIX- cyt $b_{562}(-22)/R_{10}$ | CoPPIX | 63.28± 0.54 |
| | cyt $b_{562}(-22)$ | 45.679± 0.92 |

